# IL-1-conferred gene expression pattern in ERα^+^ BCa and AR^+^ PCa cells is intrinsic to ERα^−^ BCa and AR^−^ PCa cells and promotes cell survival

**DOI:** 10.1101/773978

**Authors:** Afshan F. Nawas, Mohammed Kanchwala, Shayna E. Thomas-Jardin, Haley Dahl, Kelly Daescu, Monica Bautista, Vanessa Anunobi, Ally Wong, Rachel Meade, Ragini Mistry, Nisha Ghatwai, Felix Bayerl, Chao Xing, Nikki A. Delk

**Author notes:** Author’s contributed equally (Afshan F. Nawas, Mohammed Kanchwala). **Corresponding Author:** Nikki A. Delk^1^; 800 West Campbell Road, FO-1, Department of Biological Sciences, The University of Texas at Dallas, Richardson, Texas, USA, 75080.

## Abstract

**Background:** Breast (BCa) and prostate (PCa) cancers are hormone receptor (HR)-driven cancers. Thus, BCa and PCa patients are given therapies that reduce hormone levels or directly blocks HR activity; but most patients eventually develop treatment resistance. We have previously reported that interleukin-1 (IL-1) inflammatory cytokine downregulates *ERα* and *AR* mRNA in HR-positive (HR^+^) BCa and PCa cell lines, yet the cells can remain viable. Additionally, we identified pro-survival proteins and processes upregulated by IL-1 in HR^+^ BCa and PCa cells, that are basally high in HR^−^ BCa and PCa cells. Therefore, we hypothesize that IL-1 confers a conserved gene expression pattern in HR^+^ BCa and PCa cells that mimics conserved basal gene expression patterns in HR^−^ BCa and PCa cells to promote HR-independent survival and tumorigenicity.

**Methods:** We performed RNA sequencing (RNA-seq) for HR^+^ BCa and PCa cell lines exposed to IL-1 and for untreated HR^−^ BCa and PCa cell lines. We confirmed expression patterns of select genes by RT-qPCR and used siRNA and/or drug inhibition to silence select genes in HR^−^ BCa cell lines. Finally, we performed Ingenuity Pathway Analysis (IPA) to identify signaling pathways encoded by our RNA-seq data set.

**Results:** We identified 350 genes in common between BCa and PCa cells that are induced or repressed by IL-1 in HR^+^ cells that are, respectively, basally high or low in HR^−^ cells. Among these genes, we identified *Sequestome-1* (*SQSTM1/p62*) and *SRY* (*Sex-Determining Region Y*)*-Box 9* (*SOX9*) to be essential for survival of HR^−^ BCa and PCa cell lines. Analysis of publicly available data indicates that *p62* and *SOX9* expression are elevated in HR-independent BCa and PCa sublines generated *in vitro*, suggesting that *p62* and *SOX9* have a role in acquired treatment resistance. We also assessed HR^−^ cell line viability in response to the p62-targeting drug, verteporfin, and found that verteporfin is cytotoxic for HR^−^ cell lines.

**Conclusions:** Our 350 gene set can be used to identify novel therapeutic targets and/or biomarkers conserved among acquired (e.g. due to inflammation) or intrinsic HR-independent BCa and PCa.

## BACKGROUND

Breast (BCa) and prostate (PCa) cancer share similar etiology, where hormone receptors (HR) drive cancer cell survival^1,2^. Estrogen Receptor Alpha (ERα) promotes BCa tumor growth and Androgen Receptor (AR) promotes PCa tumor growth; thus, patients are treated with HR-targeting therapies. Unfortunately, patients can become treatment resistant due to loss of HR dependence. For example, ≥ 30% of patients that develop metastatic, castration-resistant PCa have AR-negative (AR^−^) tumors^3^ and 15-30% of BCa patients that develop endocrine resistance have tumors with reduced or lost ERα accumulation^1,4^. In addition, at the time of diagnosis, 15-20% of BCa patients are innately ERα^−^ (Triple Negative BCa)^5^ and 10-20% of PCa patients are innately AR^−^ (Small Cell Neuroendocrine PCa)^6^. Thus, there is a need to identify alternative therapeutic targets to ERα and AR hormone receptors.

Interleukin-1 (IL-1) is an inflammatory cytokine present in tumors that promotes tumor angiogenesis and metastasis^7^. IL-1 is elevated in BCa and PCa tumors^8–13^ and correlates with low or lost ERα or AR accumulation^10,14–17^. We discovered that IL-1 downregulates ERα and AR levels in HR^+^ BCa and PCa cell lines concomitant with the upregulation of pro-survival proteins that are basally high in HR^−^ cell lines^18,19^. Thus, IL-1 may select for and promote the evolution of treatment resistant cells, and our findings provide an opportunity to discover novel therapeutic targets for HR-independent BCa and PCa.

We previously used RNA sequencing (RNA-seq) to identify genes that are modulated in response to IL-1 family member, IL-1β, in the androgen-dependent AR^+^ PCa cell line, LNCaP^18^. Here, we performed RNA-seq for the estrogen-dependent ERα^+^ BCa cell line, MCF7, to identify genes that are modulated in response to both major IL-1 family members, IL-1α and IL-1β. We identified 350 genes that are conserved among IL-1-treated LNCaP and MCF7 cell lines that show similar expression patterns in untreated hormone-independent AR^−^ PC3 PCa and ERα^−^ MDA-MB-231 BCa cell lines. Not surprisingly, canonical pathway analysis reveals that the 350 gene set encodes for proteins that activate inflammatory signaling.

We selected two genes from the 350 gene set, *Sequestome-1* (*SQSTM1/p62*; hereinafter, *p62*) and *SRY* (*Sex-Determining Region Y*)-*Box 9* (*SOX9*), for functional analysis. p62 is a multi-functional scaffold protein with well-characterized roles in autophagy and antioxidant response^20^. p62 sequesters cytotoxic protein aggregates, damaged organelles, and microbes into the autophagosome for degradation and biomolecule recycling^20–26^, binds and poly-ubiquitinates Tumor Necrosis Factor Receptor-Associated Factor 6 (TRAF6), leading to the downstream activation of the pro- and anti-inflammatory transcription factor, Nuclear Factor Kappa Light Chain Enhancer of Activated B Cells (NFκB)^27,28^, and competitively binds Kelch-Like ECH-Associated Protein 1 (KEAP1) to promote activation of the antioxidant transcription factor, Nuclear Factor (Erythroid-Derived 2)-Like 2 (NRF2)^29–31^. SOX9 is a transcription factor with many diverse functions in development^32^. For example, SOX9 promotes epithelial-to-mesenchymal (EMT) transition of neural crest^33^ and endocardial endothelial^34^ cells during central nervous system and cardiac development, respectively, and induces Sertoli cell differentiation during testis development^35^. Thus, the functions of p62 and SOX9 in normal cell homeostasis and development provide cancer cells with a growth advantage and promote tumorigenicity. Specifically, we show that p62 and SOX9 are required for cell survival of HR^−^ BCa and PCa cell lines, and therefore, are rational therapeutic targets for HR-independent BCa and PCa tumors.

## METHODS

### Cell culture

MCF7 (ATCC, Manassas, VA; HTB-22)and MDA-MB-231 (ATCC, Manassas, VA; HTB-26) BCa cell lines and LNCaP (ATCC, Manassas, VA; CRL-1740), PC3 (ATCC, Manassas, VA; CRL-1435), and DU145 (ATCC, Manassas, VA; HTB-81) PCa cell lines, were grown in a 37°C, 5.0% (vol/vol) CO_2_ incubator in Dulbecco Modified Eagle Medium (DMEM; Gibco, Waltham, MA; 1185-076) supplemented with 10% FB Essence (Seradigm, Radnor, PA; 3100-500), 0.4 mM of L-glutamine (L-glut; Gibco/Invitrogen, Waltham, MA; 25030081), and 10 U/mL of penicillin G sodium and 10 mg/mL of streptomycin sulfate (pen-strep; Gibco/Invitrogen, Waltham, MA; 15140122). BT549 BCa cell line was grown in RPMI-1640 medium (Hyclone, Marlborough, MA; SH30027.01) supplemented with 10% FB essence, L-glut, and pen-strep.

### Cytokine treatment

Human recombinant IL-1α (GoldBio, St Louis MO; 1110-01A-100) and IL-1β (GoldBio, St Louis MO; 1110-01B-100) were resuspended in 0.1% bovine serum albumin (BSA, Thermo Fisher Scientific; BP 1600-1) in 1X phosphate buffered saline (PBS). Cells were treated with 25 ng/mL of IL-1 or vehicle control (0.1% BSA in 1X PBS) added to DMEM or RPMI growth medium for 5 days (BCa cell lines) or 3 days (PCa cell lines).

### RNA extraction and reverse transcription-quantitative polymerase chain reaction (RT-qPCR)

RNA was extracted from cells treated with cytokines using GeneJET RNA Purification kit (Thermo Fisher Scientific, Waltham, MA; K0732) as per manufacturer’s instructions. Genomic DNA contamination was removed by treating 1 ug of RNA with 1 U of DNase-I (Thermo scientific, Waltham, MA; EN0521) following manufacturer’s instructions. Complementary DNA was synthesized using iScript Reverse Transcription Supermix (Bio-Rad, Hercules, CA; 170-8841). RT-qPCR reactions were performed using the iTaq Universal SYBR Green Supermix (Bio-Rad, Hercules, CA; 172-5125) as per manufacturer’s instructions and the Bio-Rad CFX Connect. The cycle times for each gene was normalized to β-actin. Relative mRNA levels were calculated using 2^−ΔΔCt^ method. <u>*5′ to 3′ Primer sequences:*</u> Custom primers were obtained from Sigma-Aldrich, St. Louis, MO. *CCL20* forward, GAGTTTGCTCCTGGCTGCTTT; *CCL20* reverse, AAAGTTGCTTGCTGCTTCTGAT; *CDK2* forward, CGAGCTCCTGAAATCCTCCTG; *CDK2* reverse, GCGAGTCACCATCTCAGCAA; *CD68* forward, CAGGGAATGACTGTCCTCACA; *CD68* reverse, CAGTGCTCTCTGTAACCGTGG; *CXCR7* forward, ACGTCTGCGTCCAACAATGA; *CXCR7* reverse, AAGCCCAAGACAACGGAGAC; *IL-8* forward, ACACTGCGCCAACACAGAAAT; *IL-8* reverse, AACTTCTCCACAACCCTCTGC; *MMP16* forward, TCAGCACTGGAAGACGGTTG; *MMP16* reverse, AAATACTGCTCCGTTCCGCA; *PLK1* forward, TTCGTGTTCGTGGTGTTGGA; *PLK1* reverse, GCCAAGCACAATTTGCCGTA; *SOX9* forward, GAGACTTCTGAACGAGAGCGA; *SOX9* reverse, CGTTCTTCACCGACTTCCTCC; *Zeb1* forward, TGTACCAGAGGATGACCTGC; *Zeb1* reverse, CTTCAGGCCCCAGGATTTCTT; *p62* forward, AAATGGGTCCACCAGGAAACTGGA; *p62* reverse, TCAACTTCAATGCCCAGAGGGCTA; *β-actin* forward, GATGAGATTGGCATGGCTTT; *β-actin* reverse, CACCTTCACCGGTCCAGTTT.

### RNA sequencing (RNA-seq) analysis

RNA-seq was performed by the Genome Center at The University of Texas at Dallas (Richardson, TX). Fastq files were checked for quality using fastqc (v0.11.2)^36^ and fastq_screen (v0.4.4)^37^ and were quality trimmed using fastq-mcf (ea-utils/1.1.2-806)^38^. Trimmed fastq files were mapped to hg19 (UCSC version from igenomes) using TopHat^39^, duplicates were marked using picard-tools (v1.127 https://broadinstitute.github.io/picard/), read counts were generated using featureCounts^40^ and differential expression analysis was performed using edgeR^41^. Differential gene expression lists were generated using the following cut-offs: log_2_ counts per million (CPM) ≥ 0, log_2_ fold change (FC) ≥ 0.6 or ≤ −0.6, false discovery rate (FDR) ≤ 0.05. Pathway analysis was conducted using QIAGEN’s Ingenuity Pathway Analysis tool (http://www.qiagen.com/ingenuity). RNA-seq datasets generated for this study are available at GEO NCBI, accession GSE136420.

### Western blot and antibodies

Protein was isolated from cells using NP40 lysis buffer (0.5% NP40 [US Biological, Salem, MA; N3500], 50mM of Tris [pH 7.5], 150 mM of NaCl, 3 mM of MgCl_2_, 1X protease inhibitors [Roche, Mannheim, Germany; 05892953001]). Protein concentration was measured using the Pierce BCA Protein Assay Kit (Thermo Fisher Scientific, Waltham, MA; 23227). For Western blot analysis, equal protein concentrations were loaded onto and separated in 12% (wt/vol) sodium dodecyl sulfate polyacrylamide gel (40% acrylamide/bisacrylamide solution; Bio-Rad, Hercules, CA; 161-0148). Proteins were transferred from the gel to 0.45 μm pore size nitrocellulose membrane (Maine Manufacturing, Sanford, ME; 1215471) and total protein visualized using Ponceau S (Amresco, Radnor, PA; K793). The membrane was blocked with 2.5% (wt/vol) BSA (Thermo Fisher Scientific, Waltham, MA; BP 1600-1) in 1X tris-buffered saline with Tween 20 (TBST; 20 mM of Tris, pH 7.6, 150 mM of NaCl, 0.05% Tween-20). Primary and secondary antibodies were diluted in 2.5% BSA in 1X TBST. Protein blot bands were visualized using Clarity Western ECL Substrate (Bio-Rad, Hercules, CA; 1705061) and imaged using Amersham Imager 600 (GE, Marlborough, MA). <u>*Primary antibodies:*</u> p62 (Abcam, Cambridge, MA; H00008878-M01), SOX9 (Cell Signaling, Danvers, MA; 82630S), β-actin (Santa Cruz Biotechnology, Dallas, TX; sc-69879). <u>*Secondary antibodies:*</u> sheep anti-mouse (Jackson ImmunoResearch Laboratories, West Grove, PA; 515-035-062), goat anti-rabbit (Sigma-Aldrich, St. Louis, MO; A6154).

### Small interfering RNA (siRNA) and drug treatments

<u>*siRNA:*</u> Cells were transfected with a pool of four unique *p62* siRNA duplexes (Dharmacon, Lafayette, CO; M-010230-00-0020) or *SOX9* siRNA duplexes (Dharmacon, Lafayette, CO; M-021507-00-0010) using siTran 1.0 transfection reagent (Origene, Rockville, MD; TT300001). Non-targeting siRNA duplex was used as a negative control (Dharmacon, Lafayette, CO; D-001210-02-20). qPCR was used to confirm mRNA knock-down. <u>*drug:*</u> Verteporfin (Sigma-Aldrich, St. Louis, MO; SML0534) was resuspended in DMSO. Cells were exposed to 10 μM verteporfin or DMSO and western blotting or immunostaining for p62 oligomerization was performed to determine treatment efficacy.

### Cell counts

Cells were fixed with cold 100% methanol for 15 minutes. The nuclei were then stained with DAPI (Roche Diagnostics, Basel, Switzerland; 10236276001) and counted on the Cytation3 Imaging Reader (BioTek, Winooski, VT).

### MTT [3-(4,5-Dimethylthiazol-2-yl)-2,5-Diphenyltetrazolium Bromide] assay

MTT assay (Trevigen; 4890-25-K) was performed according to manufacturer’s instructions. Cell viability was quantified as the optical density (OD) read at a wavelength of 540 nm minus OD at 650 nm. OD was measured using the Cytation3 Imaging Reader (BioTek, Winooski, VT).

### Immunostaining

Cells were fixed and permeabilized with 100% methanol at −20°C for 30 minutes. Fixed cells were blocked with 2.5% BSA in 1X PBS at room temperature for at least 30 minutes. Antibodies were diluted in 2.5% BSA in 1X PBS. Cells were incubated in primary antibody overnight at 4°C, washed with 1X PBS, and then incubated with fluorescently labeled secondary antibody overnight at 4°C in the dark. Primary antibody: p62 (Santa Cruz Biotechnology, Dallas, TX; sc-28359). Fluorescently labeled secondary antibody: Alexa Fluor 488, goat anti-mouse (Invitrogen, Waltham, MA; A11001). Nuclei were stained with DAPI (Roche Diagnostics, Basel, Switzerland; 10236276001). Immunostained cells were imaged at 40X magnification using a Nikon epifluorescence microscope (Nikon, Melville, NY).

### Statistical analysis

Statistical significance was determined using unpaired student t-test. P-value ≤ 0.05 is considered statistically significant. Graphs are shown as the average of a minimum of n = 3 biological replicates ± standard deviation (STDEV).

## RESULTS

### Identification of IL-1 conferred gene signature in hormone receptor positive BCa and PCa cell lines that mimic basal gene expression pattern in hormone receptor negative BCa and PCa cells

We previously found that IL-1 represses hormone receptors in ERα^+^/PR^+^ BCa^19^ and AR^+^ PCa^18,42^ cell lines concomitant with p62 upregulation, while ERα^−^/PR^−^ BCa and AR^−^ PCa cell lines intrinsically have high basal p62. This led us to speculate that IL-1 elicits similar changes in gene expression in hormone receptor positive (HR^+^) BCa and PCa cells that mimic intrinsic gene expression patterns in hormone receptor negative (HR^−^) BCa and PCa cells. Such changes would enable BCa and PCa cells to elicit compensatory survival pathways in the absence of hormone receptor activity and, thus, these changes in gene expression could confer resistance to hormone therapy for BCa and PCa tumor cells. To identify the conserved gene signature conferred by IL-1 in HR^+^ BCa and PCa cells that mimics intrinsic gene expression patterns in HR^−^ BCa and PCa cells, we performed RNA sequencing (RNA-seq) followed by differential gene expression analysis (log_2_ CPM ≥ 0, log_2_ FC ≥ 0.6 or ≤ −0.6, and FDR ≤ 0.05) for IL-1α- or IL-1β-treated ERα^+^/PR^+^ MCF7 BCa cell line, IL-1β-treated AR^+^ LNCaP PCa cell line, vehicle control-treated ERα^−^/PR^−^ MDA-MB-231 BCa cell line, and vehicle control-treated AR^−^ PC3 PCa cell line. LNCaP and PC3 RNA-seq data was previously reported^18^ (GSE105088). Five sets of differential gene expression analysis were performed: (Set 1) MCF7 cells treated with vehicle control versus IL-1α (“MCF7_VEH-VS-MCF7_IL1A”); (Set 2) MCF7 cells treated with vehicle control versus IL-1β (“MCF7_VEH-VS-MCF7_IL1B”); (Set 3) MCF7 cells treated with vehicle control versus MDA-MB-231 cells treated with vehicle control (“MCF7_VEH-VS-231_VEH”); (Set 4) LNCaP cells treated with vehicle control versus IL-1β (“LNCaP_VEH-VS-LNCaP_IL1B”); and LNCaP cells treated with vehicle control versus PC3 cells treated with vehicle control (“LNCaP_VEH-VS-PC3_VEH”) (Fig. 1A, Supplemental Table 1). We identified 2,735 genes in the intersection of Set 1, 2, and 3 and of those genes, 1,707 are consistent in fold change direction (Set 6). Set 6 are the genes that are induced or repressed by both IL-1α and IL-1β in MCF7 cells that are, respectively, basally high or low in MDA-MB-231 cells. We identified 2,786 genes in the intersection of Set 4 and 5 and of those genes, 1,900 are consistent in fold change direction (Set 7). Set 7 are the genes that are induced or repressed by IL-1β in LNCaP cells that are, respectively, basally high or low in PC3 cells. Finally, we identified the 420 genes in the intersection of Set 6 and 7 and of those genes, 350 are consistent in fold change direction (Set 8). Set 8 are the genes that are induced or repressed by IL-1 in MCF7 and LNCaP cells that are, respectively, basally high or low in MDA-MB-231 and PC3 cells. Thus, the 350 gene set represents conserved genes in both BCa and PCa cells that are expected to promote cell survival and tumorigenicity when hormone receptor signaling is lost.

**Figure 1.**
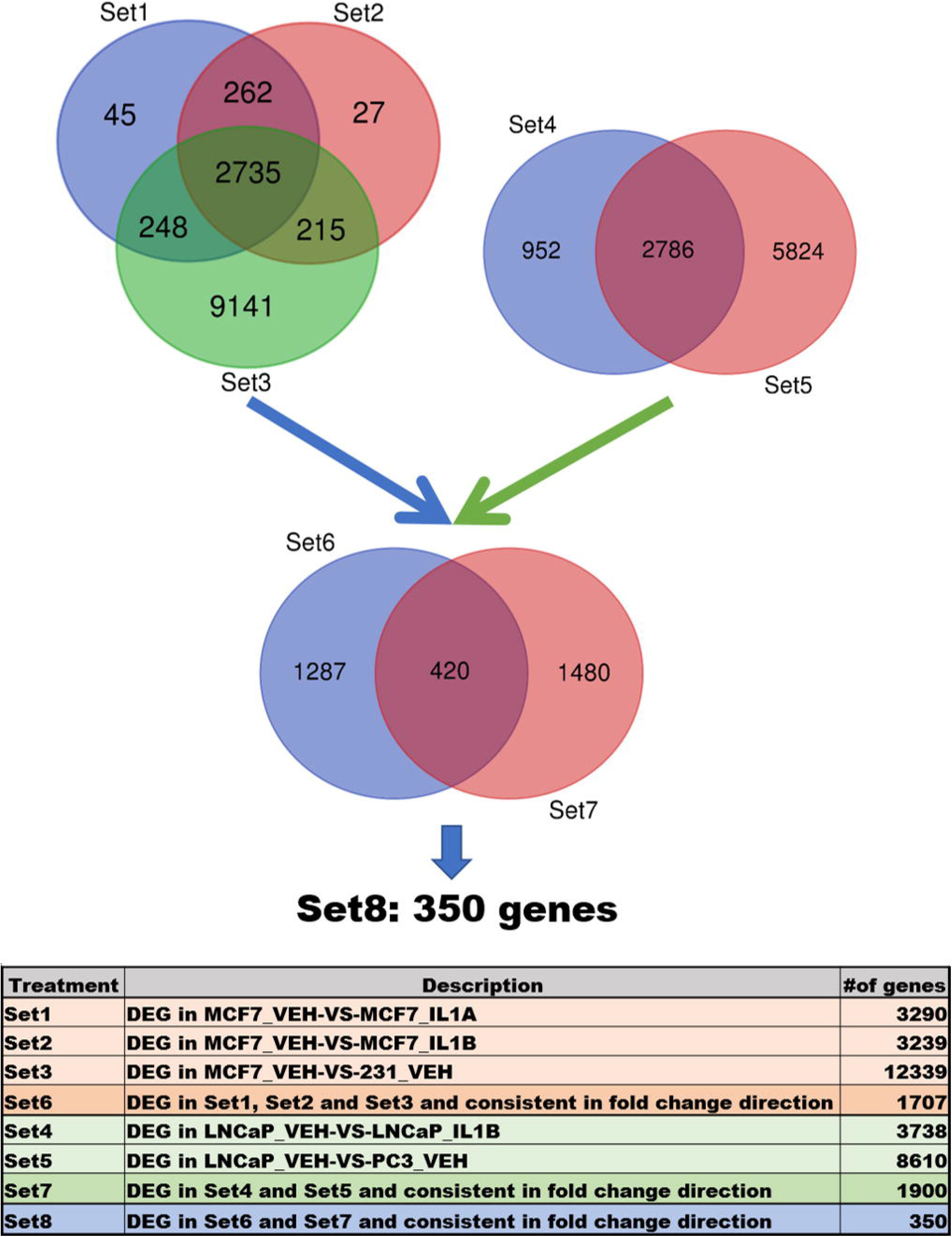
Workflow of data overlap to find 350 gene signature. Sets 1, 2, 3, 4 and 5 are generated from the RNA-seq data using following cut-offs: log_2_ CPM ≥ 0, absolute log_2_ FC > 0.6, and FDR ≤ 0.05. Sets 6, 7 and 8 are generated as described in the table. DEG = Differentially Expressed Genes; CPM = Counts Per Millions; FC = Fold Change; FDR = False Discovery Rate.

### Validation of select genes from 350 gene signature

We selected six upregulated genes (*CCL20*, *CD68*, *IL-8*, *p62*, *SOX9*, *Zeb1*) and four downregulated genes (*CDK2*, *CXCR7*, *MMP16*, *PLK1*) from our 350 gene set to validate by RT-qPCR. MCF7 and LNCaP cells were treated with 25 ng/ml IL-1α or IL-1β and MDA-MB-231 and PC3 cells were treated with vehicle control. RT-qPCR confirmed that IL-1 induces *CCL20*, *IL-8*, *p62*, and *Zeb1* in HR^+^ MCF7 (Fig. 2A) and LNCaP (Fig. 2B) cells. We detected a significant increase in *CD68* and *SOX9* mRNA (Fig. 2B) and/or protein (Fig. 3A) in IL-1-treated LNCaP cells. We did not detect an increase in *SOX9* mRNA in IL-1-treated MCF7 cells (Fig. 2A) and only a slight increase in SOX9 protein (Fig. 3A); and *CD68* mRNA levels were only slightly induced by IL-1β in MCF7 cells (Fig. 2A). Finally, RT-qPCR confirmed that IL-1α and IL-1β repress *CDK2*, *CXCR7*, *MMP16*, and *PLK1* in MCF7 and LNCaP cells (Fig. 2).

**Figure 2.**
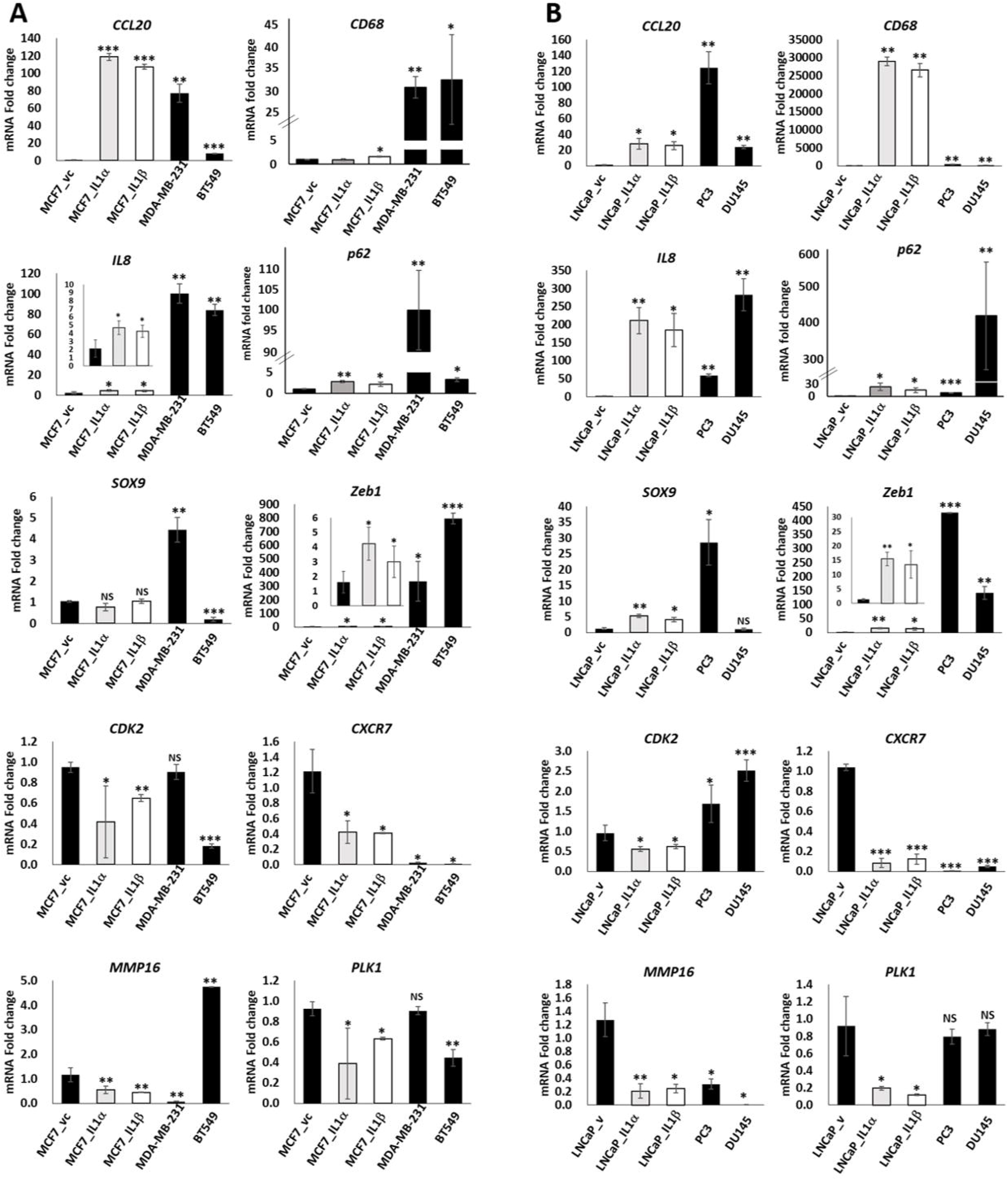
RT-qPCR validation of select genes from 350 gene signature. RT-qPCR was performed for select genes for MCF7 and LNCaP cell lines treated with vehicle control or 25 ng/ml IL-1α or IL-1β for 5 days. MDA-MB-231, BT549, PC3 and DU145 cell lines were treated with vehicle control only. (A) RT-qPCR shows that *CCL20*, *CD68*, *IL8*, *p62*, *SOX9* and *Zeb1* are induced by IL-1 in MCF7 and/or LNCaP cell lines and are basally high in MDA-MB-231, BT549, PC3 and/or DU145 cell lines. (B) RT-qPCR shows that *CXCR7* and *MMP16*, but not *CDK2* or *PLK1*, are downregulated by IL-1 in MCF7 and/or LNCaP cell lines and are basally high in MDA-MB-231, BT549, PC3 and/or DU145 cell lines. N = 3 biological replicates; error bars, +/−STDEV; p-value, * ≤ 0.05, ** ≤ 0.005, ***≤ 0.0005. mRNA fold change is normalized to MCF7 vehicle control for the BCa cell lines and to LNCaP vehicle control for the PCa cell lines.

**Figure. 3.**
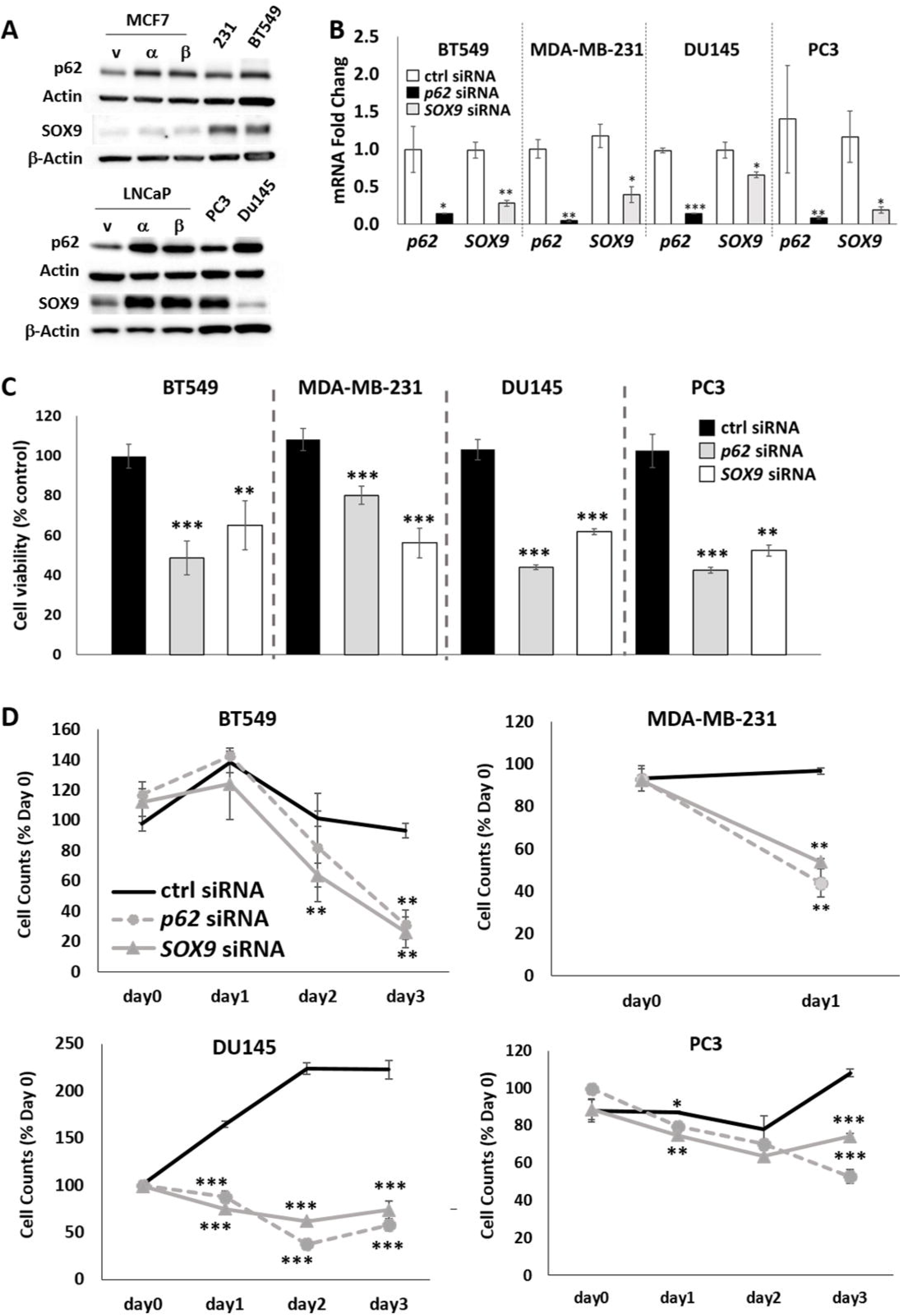
p62 and SOX9 are cytoprotective for hormone receptor negative BCa and PCa cell lines. (A) Western blot analysis was performed for MCF7 or LNCaP cells treated with vehicle control or 25 ng/ml IL-1 or for untreated MDA-MB-231, BT549, PC3, and DU145 cells. p62 and SOX9 are induced by IL-1 in MCF7 and LNCaP cells and p62 and/or SOX9 protein is basally high in MDA-MB-231, BT549, PC3, and DU145 cells. MDA-MB-231, BT549, PC3, and DU145 cell lines were treated with 70 nM control siRNA, *p62* siRNA, or *SOX9* siRNA and (B) after 1 day in siRNA, RT-qPCR was performed to validate *p62* or *SOX9* knock-down, (C) MTT was performed after 1 day (MDA-MB-231) or 3 days (BT549, PC3, DU145) in siRNA, or (D) cell counts were recorded at day 0 (no treatment), day 1, 2, and 3. Loss of p62 or SOX9 is cytotoxic for the MDA-MB-231, BT549, PC3, and DU145 cell lines. N = 3-4 biological replicates; error bars, +/−STDEV; p-value, * ≤ 0.05, ** ≤ 0.005, ***≤ 0.0005. mRNA fold change is normalized within each cell line to control siRNA.

In addition to MDA-MB-231 and PC3 cell lines, we performed RT-qPCR for basal gene expression in an additional ERα^−^/PR^−^BCa cell line, BT549, and an additional AR^−^ PCa cell line, DU145. RT-qPCR confirmed that *CCL20*, *CD68*, *IL-8*, *p62*, and *Zeb1* are basally high MDA-MB-231, BT549, PC3, and DU145 cells (Fig. 2). *SOX9* mRNA (Fig. 2) and protein (Fig. 3A) are basally high in MDA-MB-231 and PC3 cells and SOX9 protein is basally high in BT549 (Fig. 3A); however, *SOX9* mRNA is not basally high in BT549 or DU145 (Fig. 2). We did not find that the select IL-1-repressed genes are consistently downregulated in the HR^−^ BCa and PCa cell lines (Fig 2). Taken together, RT-qPCR primarily confirmed the 350 gene set expression patterns identified in MCF7, LNCaP, MDA-MB-231, and PC3 cells by RNA-seq.

### p62 and SOX9 are cytoprotective for HR^−^ BCa cell lines

Given that IL-1 reduces hormone receptors concomitant with the upregulation of pro-survival and pro-tumorigenic genes, such as p62^42–48^, we hypothesized that IL-1 can promote resistance to hormone receptor-targeted therapy. We compared our 350 gene set to published RNA-seq data from enzalutamide-resistant LNCaP^3^ (GSE99381; APIPC subline versus APIPC_P (parental)) or fulvestrant-resistant MCF7^49^ (GSE74391; ICI182R1 or ICI182R6 subline versus fulvestrant-sensitive subline (parental)) sublines. Among the select upregulated genes we chose for RT-qPCR confirmation, *p62* and *SOX9* were upregulated in both treatment-resistant subline models (Supplemental Table 1). Downregulation of *AR* or *ERα/PR* and target gene expression in the LNCaP^3^ or MCF7^49^ treatment-resistant sublines (Supplemental Table 1) indicates these models evolved to survive without hormone receptor activity. Therefore, we siRNA-silenced *p62* and *SOX9* in HR^−^ BCa and PCa cell lines to determine if p62 or SOX9 are required for viability in cells that intrinsically lack hormone receptor activity. MDA-MB-231, BT549, PC3, and DU145 cells were transfected with *p62* or *SOX9* siRNA (Fig. 3B) and cell viability was determined on day 1 or 3 using MTT (Fig. 3C) or by recording cell number on day 1, 2, and 3 (Fig. 3D). *p62* and *SOX9* siRNA are cytotoxic for the HR^−^ BCa and PCa cell lines. Thus, p62 and SOX9 promote cell survival for cells that intrinsically lack hormone receptor activity.

### Verteporfin is cytotoxic for HR^−^ BCa and PCa cell lines

Cell that intrinsically lack hormone receptor activity are not susceptible to hormone receptor-targeting drugs, such as enzalutamide or fulvestrant. Therefore, alternative therapeutic targets are needed. The p62 inhibitor, verteporfin (Visudyne®), is an FDA-approved, photosensitizing drug used with laser light to treat leaky blood vessels in the eye caused by macular degeneration. Currently, verteporfin is being tested in a Phase I clinical trial as a photosensitizer for the SpectraCure P18 photodynamic therapy system for recurrent PCa (NCT03067051). Recently, verteporfin, alone, was shown to reduce the tumor growth of subcutaneous prostate epithelial xenografts overexpressing *p62* and verteporfin was able to reduce the growth of PC3 xenografts^43^. Verteporfin oligomerizes p62 (Fig. 4A & B), thereby preventing p62 interaction with binding partners and inhibiting p62 function^43,50^. We treated HR^−^ BCa and PCa cell lines with 10 μM verteporfin for 5 days and assayed cell viability using MTT (Fig. 4C) or by recording cell number on day 1, 3, and 5 (Fig. 4D). Verteporfin is cytotoxic for HR^−^ BCa and PCa cell lines.

**Figure. 4.**
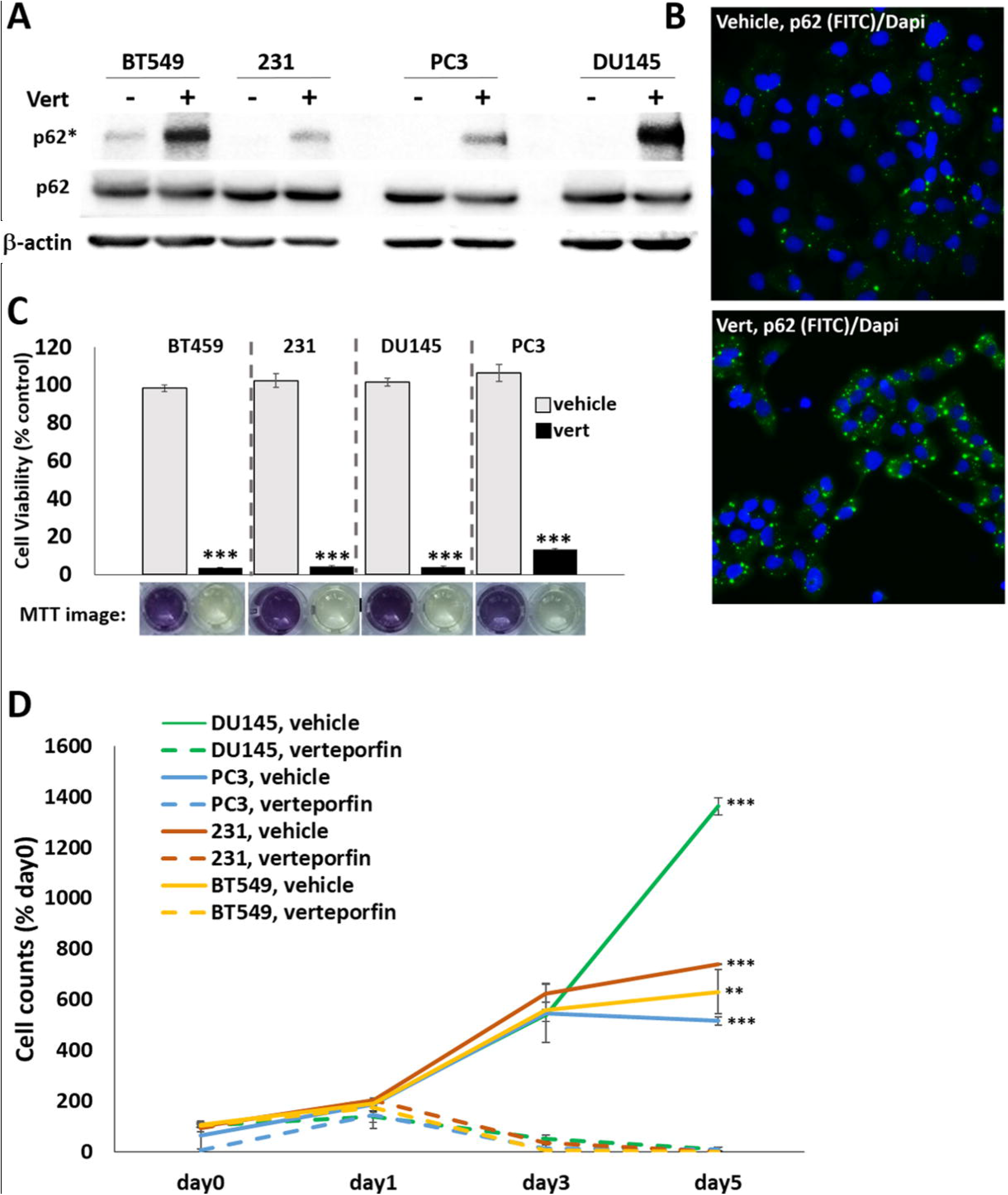
Verteporfin is cytotoxic for HR^−^ BCa and PCa cell lines. (A) MDA-MB-231, BT549, PC3, and DU145 cell lines were treated with vehicle control or 10 μM verteporfin for 1 day. Western blot analysis shows oligomerized p62 (p62*), indicating treatment efficacy. (B) Representative images of DU145 cells treated with vehicle control or 5 μM verteporfin for 7 days and immunostained for p62 (FITC; nuclei, DAPI; 40X magnification). Verteporfin induces p62 oligomerization indicated by p62 puncta. Cells were treated with vehicle control or 10 μM verteporfin for 5 days and (C) MTT assay was performed to assess cell viability or (D) cell counts were taken at day 0 (no treatment), day 1, 3, and 5. Verteporfin is cytotoxic for MDA-MB-231, BT549, PC3, and DU145 cell lines. N = 3 biological replicates; error bars, +/−STDEV; p-value, ** ≤ 0.005, ***≤ 0.0005.

### The 350 gene signature maps to pro-tumorigenic processes in BCa and PCa cells

We used the IPA Canonical Pathways module to identify signaling pathways represented in our 350 gene set and found that the gene set encodes for proteins that are predicted to activate inflammatory signaling, including interferon (z-score = 3.2, −log p-value = 9.86E+00), IL-1 (z-score = 2, −log p-value 1.26E+00), and IL-8 signaling (z-score = 1.7, −log p-value = 4.10E+00) (Supplemental Table 1). We also used the IPA Regulator Effects module to predict cancer-specific networks encoded by our 350 gene set and we selected the networks in which both p62 and SOX9 were target molecules. We found three such networks in which upstream regulators CTNNB1 (z-score 2.738, p-value 5.12E-07), FGF2 (z-score 2.091, p-value 2.21E-09) and TNF (z-score 7.192, p-value 4.87E-28) were found to be activated and predicted to promote neoplasia of cells (z-score 2.699, p-value 5.20E-12 (CTNNB1)) and malignancy (z-score 2.251, p-value 1.41E-13 (FGF2, TNF)) (Supplemental Table 1). Thus, the IL-1-conferred 350 gene set encodes multiple different pro-tumorigenic signaling pathways conserved among multiple different regulators.

### Predicted p62 and SOX9 target molecules and functional networks in the 350 gene signature

Finally, given that siRNA-mediated loss of *p62* or *SOX9* are cytotoxic for HR^−^ BCa and PCa cells, we used the IPA Upstream Regulators module to identify p62 or SOX9 predicted target molecules and functional networks in the 350 gene data set. p62 is predicted to induce *CXCL2, IL15RA, IRF1, PLAT, RGCC*, and *RSAD2* expression (z-score = 2.449, p-value = 8.38E-05) (Fig. 5A; Supplemental Table 1) and predicted to activate HIF1A, IL-1β, NFκB1, NFKBIA, NFκB (complex), NR3C1, RELA, and SQSTM1(p62) signaling (Fig. 5B; Supplemental Table 1). The p62-regulated genes and interactive networks are known mediators of inflammatory signaling and immunity (CXCL2, IL15RA, IRF1, RSAD2, IL-1β, NFκB, NR3C1, RELA, SQSTM1(p62)), hypoxia (HIF1A), fibrinolysis (PLAT), and cell cycle regulation (RGCC). SOX9 did not appear as a master regulator in IPA; therefore, we manually extracted its interactome from IPA and overlapped the results with the 350 gene list to identify relevant interactions. SOX9 is predicted to activate *VNN1* expression and mediate FN1 and T-Cell Factor (TCF) signaling (Fig. 5C). VNN1 functions in inflammation and immunity, FN1 promotes wound healing, and TCF interacts with β-catenin to mediate WNT signaling. Disruption of any of the processes that p62 or SOX9 are predicted to regulate or mediate as part of the 350 gene signature would be expected to reduce cell viability and may explain *p62* or *SOX9* siRNA-mediated cytotoxicity in BCa and PCa cells.

**Figure 5.**
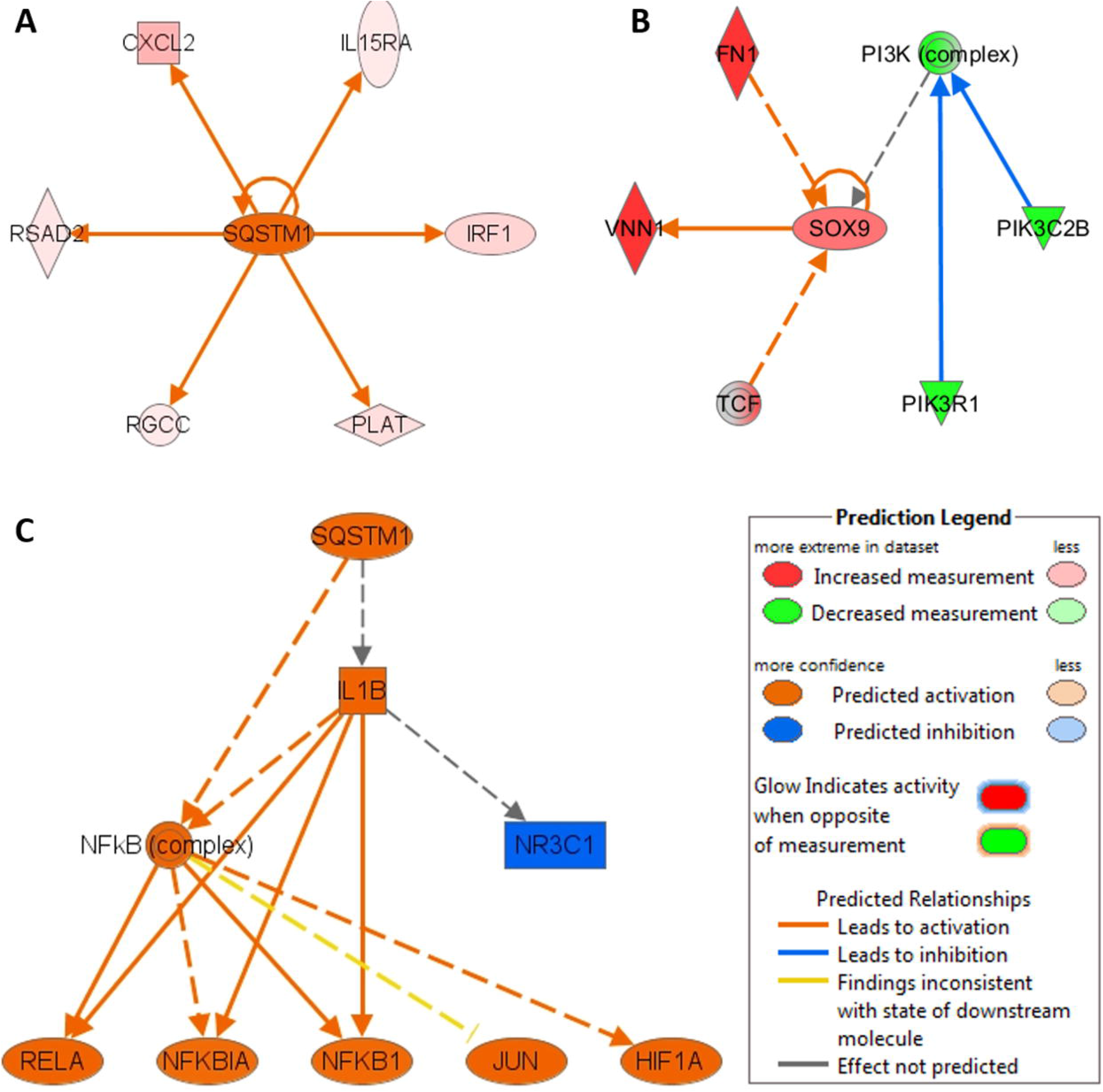
Regulatory Networks generated using IPA and 350 gene signature. (A) Interactome of p62 identified using Upstream Regulator Analysis of IPA (z-score = 2.449, p-value = 8.38E-05). (B) Interactome of SOX9 identified using IPA (IPA does not report SOX9 as an upstream regulator). (C) Mechanistic Network identified using IPA where p62 is a master regulator (bias-corrected p-value = 8.00E-03).

## DISCUSSION

### IL-1 conferred 350-gene signature is enriched in genes and pathways that promote BCa and PCa survival and progression

We have previously shown that HR^low/−^ BCa^19^ and PCa^18,42^ cells are enriched when HR^+^ cells are exposed to IL-1. Here, we sought to compare the gene expression pattern overlap between cells that lose hormone receptors in response to IL-1 to cells that are intrinsically HR^−^. We identified 1707 genes in BCa cells and 1900 genes in PCa cells that are upregulated by IL-1 in HR^+^ cells and basally high in HR^−^ cells or downregulated by IL-1 in HR^+^ cells and basally low in HR^−^ cells. To identify genes that were common to both BCa and PCa, we looked for overlapping genes and filtered down to a set of 350 genes. This conserved expression pattern represents IL-1-induced signaling in HR^+^ BCa and PCa cells and constitutive signaling in HR^−^ BCa and PCa cells. To gain insight in the functional output of the 350 gene set, we used IPA to predict canonical pathways. IL-1 is an inflammatory cytokine, therefore, as expected, the 350 gene set showed an enrichment for inflammatory pathways like IL-1, IL-8, IL-6, interferon, and TNF receptor signaling. Importantly, inflammatory signaling promotes BCa^9^ and PCa^51^ progression.

We used IPA to identify cancer-specific networks encoded by the 350 gene set that also include p62 and SOX9 as downstream target signaling molecules and found that IL-1 is predicted to activate molecular programs similar to CTNNB1, Fibroblast Growth Factor (FGF2), and Tumor Necrosis Factor (TNF). *CTNNB1* encodes for beta-catenin (β-catenin) and mediates the canonical WNT pathway regulation of cell fate and tissue development^52^. WNT-β-catenin signaling promotes tumor progression, stem cell proliferation and treatment resistance in BCa^53^ and PCa^54^. FGF2 signaling regulates normal tissue development, angiogenesis, and wound healing^55^ and FGF2 promotes tumor progression, metastasis, and/or angiogenesis in BCa^56^ and PCa^57^. Finally, TNF is an inflammatory cytokine that promotes BCa EMT, metastasis, and invasion^58^ and promotes PCa androgen independence, EMT, metastasis, and invasion^59^. Taken together, the IL-1-conferred 350 gene signature encodes for proteins that have been shown to participated in conserved pro-tumorigenic pathways in response to multiple different stimuli.

### p62 and SOX9 are rational therapeutic targets for BCa and PCa when hormone receptor signaling is lost

p62 protein is overexpressed in PCa patient tumors, is prognostic, and correlates with advanced PCa disease^60,61^ and p62 protein accumulation is elevated in BCa patient tumors relative to the normal adjacent tissue^47^. Furthermore, SOX9 was found to be highly expressed in BCa patient tumors relative to normal tissue and showed strongest expression in ERα^−^ tumors^62^ and elevated SOX9 levels correlates with disease progression in PCa^63^. Analysis of publicly available gene expression data from ERα^−^, fulvestrant-resistant MCF7 and AR^−^, enzalutamide-resistant LNCaP sublines showed that *p62* and *SOX9* are upregulated. Fulvestrant and enzalutamide, respectively, block activity of ERα and AR, thus HR^−^ BCa and PCa are intrinsically resistant to these HR-targeting drugs. We found that gene silencing *p62* or *SOX9* in HR^−^ BCa and PCa cell lines is cytotoxic, including treatment with the p62-targeting drug, verteporfin. Taken together, cells that evolve HR-independence and treatment resistance could concomitantly evolve dependency on cytoprotective, pro-tumorigenic p62 or SOX9, making p62 and SOX9 rational therapeutic targets in BCa and PCa.

IPA predicts p62 to be a master regulator of inflammation, immunity, hypoxia, fibrinolysis, and the cell cycle and predicts SOX9 to function in immunity and WNT signaling through their respective interactions within the 350 gene set. For example, p62-NRF2 signaling was shown to attenuate reactive oxygen, maintain stemness and promote tumor growth of BCa cells^64^ and SOX9 was shown to activate WNT signaling and drive WNT-mediated PCa tumor growth^65^. Thus, the IPA-predicted cellular functions of p62 and SOX9 in the context of the 350 gene set provides insight into how p62 and SOX9 promote BCa and PCa survival and, in particular, promote survival when HR signaling is lost. Equally important, in addition to p62 and SOX9, our 350 gene set provides a myriad of additional rational target molecules and networks conserved among both treatment-resistant BCa and PCa and, thus, our 350 gene set has the potential to have a broader patient impact.

## CONCLUSION

Having discovered that IL-1 represses hormone receptors in both ERα^+^ BCa and AR^+^ PCa cell lines, we identified a conserved gene expression signature between IL-1-treated HR^+^ BCa and PCa cell lines and HR^−^ cell lines. We performed functional bioinformatics analyses to predict cellular function of the gene signature. This approach revealed potential therapeutic targets, p62 and SOX9, and identified signaling pathways downstream of inflammatory (e.g. IL-1 and TNF), FGF, or WNT signaling that could be targeted in both treatment-resistant BCa and PCa.

## LIST OF ABBREVIATIONS

AR: Androgen Receptor
CTNNB1: Beta Catenin
BCa: Breast Cancer
CXCL2: C-X-C Motif Chemokine Ligand 2
EMT: epithelial-to-mesenchymal transition
ERα: Estrogen Receptor Alpha
FGF2: Fibroblast Growth Factor 2
FN1: Fibronectin 1
HIF1A: Hypoxia Inducible Factor 1 Subunit Alpha
HR: Hormone Receptor
IL15RA: Interleukin 15 Receptor Subunit Alpha
IL-1a: Interleukin-1 alpha
IL-1b: Interleukin-1 beta
IL-8: Interleukin-8
IPA: Ingenuity Pathway Analysis
IRF1: Interferon Regulatory Factor 1
KEAP1: Kelch-Like ECH-Associated Protein 1
NFKB: Nuclear Factor Kappa Light Chain Enhancer of Activated B Cells
NFKBIA: NFKB Inhibitor Alpha
NR3C1: Glucocorticoid Receptor
NRF2: Nuclear Factor (Erythroid-Derived 2)-Like 2
PCa: Prostate Cancer
PLAT: Tissue-Type Plasminogen Activator
RELA: NFKB Transcription Factor P65
RGCC: Regulator Of Cell Cycle
RSAD2: Radical S-Adenosyl Methionine Domain Containing 2
SOX9: SRY (Sex-Determining Region Y)-Box 9
Sqstm1: Sequestome-1
TCF: T-cell Factor
TNF: Tumor Necrosis Factor
TRAF6: Tumor Necrosis Factor Receptor-Associated Factor 6
VNN1: Vanin 1

## DECLARATIONS

### Ethics approval and consent to participate

Not applicable

### Consent for publication

Not applicable

### Availability of data and materials

RNA-seq datasets generated for this study are available at GEO NCBI, accession GSE136420.

### Authors’ contributions

AF, experiment design, execution, analysis, manuscript preparation. MK, bioinformatics analysis, manuscript preparation. STJ, optimization of PCa culture, RT-qpcr, western blot analysis, manuscript editing. HD, assisted with siRNA and cell viability experiments. KD and FB, analysis of publicly available expression data. MB, assisted with verteporfin treatments. VA, AW, R Meade, undergraduate student researchers assisted AF with experiments. R Mistry and NG, optimized PCa and BCa cell culture and treatment conditions for siRNA and cell viability. CX, head of bioinformatics core. ND, senior and corresponding author. All authors read and approved the final manuscript.

## Acknowledgments

We would like to thank the members of the Delk and Xing labs and the University of Texas at Dallas Genome Center for services in support our research.

---

**Supplemental Table 1: Gene expression and IPA data.** The “Table Legend” tab is a description of each worksheet tab. Each worksheet tab contains the gene expression data or IPA analysis described in the paper.

